# Integrating Artificial Intelligence-Driven Digital Pathology and Genomics to Establish Patient-Derived Organoids as a Novel Alternative Model for Drug Response in Head and Neck Cancer

**DOI:** 10.1101/2025.06.24.660824

**Authors:** Rose Doerfler, Jie Chen, Carl Kim, Joshua D. Smith, Micah Harris, Krishna B. Singh, Brian Isett, Rebekah E. Dadey, Daniel D. Brown, Adrian V. Lee, Xuefeng Wang, Matthew E. Spector, Seungwon Kim, Shaum Sridharan, Kevin Contrera, Katelyn Smith, Carly Reeder, Maureen Lyons, Jianhua Luo, Silvia Liu, Dan P. Zandberg, Heath D. Skinner, Ioannis K. Zervantonakis, Lazar Vujanović, Robert L. Ferris, Raja R. Seethala, José P. Zevallos, Jason J. Luke, Riyue Bao

**Affiliations:** Hillman Cancer Center, UPMC, Pittsburgh, PA, USA; Department of Medicine, University of Pittsburgh, Pittsburgh, PA, USA; Cancer Bioinformatics Core, UPMC, Pittsburgh, PA, USA; Department of Otolaryngology, University of Pittsburgh, Pittsburgh, PA, USA; Institute for Precision Medicine, UPMC, Pittsburgh, PA, USA; Department of Pharmacology and Chemical Biology, University of Pittsburgh, Pittsburgh, PA, USA; Biostatistics and Bioinformatics Program and Immuno-Oncology Program, Moffitt Cancer Center, Tampa, FL; Department of Immunology, University of Pittsburgh, Pittsburgh, PA, USA; Department of Pathology, University of Pittsburgh School of Medicine, Pittsburgh, PA, USA; Organ Pathobiology and Therapeutics Institute, University of Pittsburgh School of Medicine, Pittsburgh, PA, USA; Department of Radiation Oncology, University of Pittsburgh School of Medicine, Pittsburgh, PA, USA; Department of Bioengineering, University of Pittsburgh, Pittsburgh, PA, USA; UNC Lineberger Comprehensive Cancer Center, Chapel Hill, NC, USA; Division of Hematology/Oncology, University of Pittsburgh, Pittsburgh, PA, USA

**Keywords:** Head and neck cancer, Organoids, Novel alternative model, Drug response, Artificial Intelligence, Genomics, Digital pathology, Human specimens

## Abstract

Patient-derived organoids (PDOs) are emerging as advanced 3D *ex vivo* novel alternative method (NAM) preclinical models, offering significant advantages over traditional cell lines and monolayer cultures for therapeutic development. In this study, we established PDOs from surgically resected fresh tissues of human papillomavirus (HPV)-negative head and neck squamous cell carcinoma (HNSCC) across anatomical sites, tumor T-categories, and sample types. These PDOs faithfully recapitulate the tumor’s pathology, mutational profile, and drug response. To enable rapid classification of PDO identity, we developed a new convolutional neural network (CNN) model, TransferNet-PDO, which accurately distinguished tumor *versus* normal PDOs in culture using digital histopathology images (AUC≥0.88). PDOs maintained stable cultures and were cryopreserved between passages 5 and 12. Immunohistochemistry (IHC) staining (PanCK, p63, Cytokeratin 13, Ki67) confirmed squamous phenotype and histologic aggression of the original tumor. For tumors harboring *TP53* mutations by whole-exome sequencing (WES), PDOs retained the corresponding p53 functional status as confirmed by IHC (enhanced or loss of expression). Somatic mutational landscape revealed that PDOs preserved driver somatic mutations, copy number variations (CNVs), and clonal architecture including low-prevalence subclones. Drug sensitivity assessment of PDOs showed that cisplatin reduced cell viability, whereas cetuximab and lenvatinib had minimal effects. Chemoradiation led to greater tumor organoid killing compared to radiation or chemotherapy alone. This study presents an integrated HNSCC PDO platform combining tissue biobanking, organoid establishment, multi-omics characterization, functional drug screening, and AI-driven histopathologic classification, providing a comprehensive and scalable system for translational cancer research.

## Introduction

Head and neck squamous cell carcinoma (HNSCC) remains a significant clinical challenge, especially in carcinogen-driven, Human Papillomavirus (HPV)-negative HNSCC populations^1^. Despite intensive standard therapies, 40%∼60% of patients experience disease progression or die within five years. There is an urgent need to improve therapeutic strategies in both the curative intent locally advanced setting and the recurrent/metastatic setting^2 3^.

Historically, murine models have been widely used in HNSCC research, providing critical in vivo insights into tumor initiation, progression, and metastasis^4^. However, the translational success remains limited due to fundamental differences between murine and human biology, high costs, lengthy timelines, lack of reproducibility, and low scalability^5^. These challenges have driven efforts in developing novel alternative models (NAMs) to better recapitulate human biology. Among these, patient-derived organoids (PDOs), three-dimensional (3D) cultures derived from patient tumors, emerge as highly promising for their scalability, genomic fidelity, and translational potential in personalized therapy development^6^.

While PDOs have been well established in cancers such as colorectal^7 8^, lung^9 10^, and brain^11 12^, models for HNSCC remain relatively underdeveloped. Global efforts are increasing^13–22^ however challenges persist including overgrowth of normal epithelial or stromal cells, difficulty preserving tumor heterogeneity and clonality over passages, and limited access to fresh surgical specimens. A particularly underexplored area is the ability to accurately distinguish malignant from non-malignant organoids within cultures, which is essential for translational applications. Artificial intelligence (AI), including deep convolutional neural networks (CNNs), has transformed pathology by enabling rapid, quantitative, and precise malignancy detection in digital histopathology^23^. While PDOs recapitulate tumor morphology and genomics, their in vitro growth patterns could present unique challenges for image interpretation^24^. In this context, AI-driven classifiers offer key advantages by detecting subtle single-cell morphologic features, such as nuclear eccentricity or irregular cell borders, across large datasets^25–28^. However, most existing pathology AI tools were trained on primary human tissues^29–35^, and their application to organoid systems remains largely unexplored. To our knowledge, no AI-based malignancy classifier has been developed specifically for HNSCC PDOs. Integrating AI into this space therefore presents a timely opportunity to enhance the accuracy, scalability, and translational utility of organoid models in HNSCC.

In this study, we directly address these challenges by establishing a robust cohort of PDOs from fresh HPV-negative HNSCC tumors in patients across diverse clinical characteristics (**Fig. 1A**). We introduce TransferNet-PDO, a single-cell CNN classifier that accurately distinguishes tumor from normal PDOs using histopathology images. We further show that the clonal architecture of the original tumor is preserved in serial PDO passages, including low-frequency subclones. Finally, we demonstrate clinically relevant drug sensitivity to standard and targeted therapies, highlighting the value of PDOs in preclinical therapeutic testing and precision oncology. Our work advances PDO development and lays the foundation for personalized treatment strategies in HNSCC.

**Figure 1.**
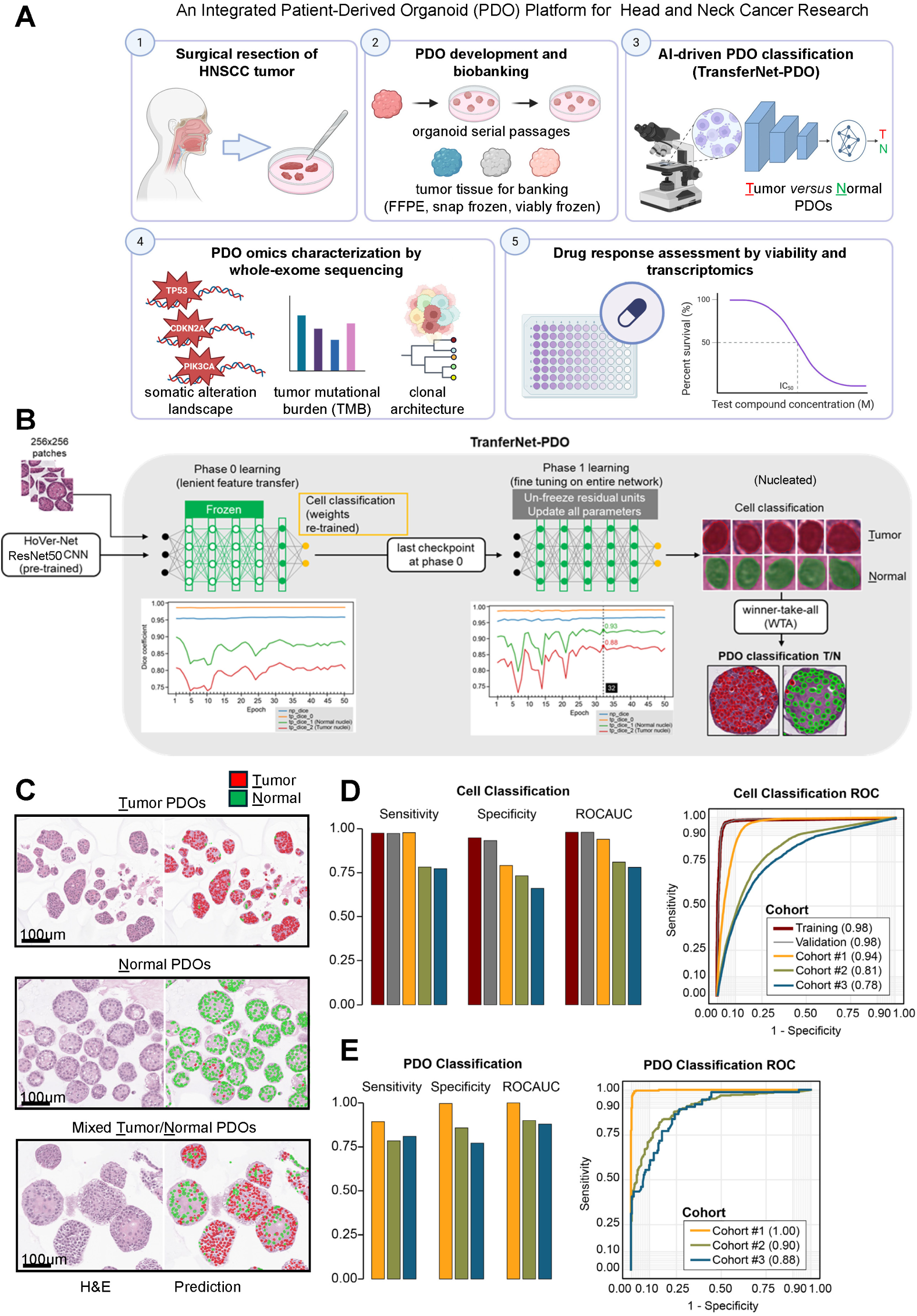
TransferNet-PDO accurately distinguishes malignant from normal PDOs in HNSCC. (**A**) Study design for PDO development and analysis. Five main modules are shown: (1) Fresh tumor specimens from patients with HNSCC are collected at the time of surgical resection; (2) PDOs are developed and expanded through serial passaging; tissue aliquots are banked using multiple preservation methods (FFPE, snap frozen, viably frozen) when available; (3) A new CNN classifier (TransferNet-PDO) is developed to distinguish tumor *versus* normal PDOs from H&E images; (4) PDOs undergo whole-exome sequencing to characterize somatic alterations, tumor mutational burden (TMB), and clonal architecture in comparison to the original tumor; (5) Drug sensitivity is assessed by cell viability assays and transcriptomic analysis. (**B**) ransferNet-PDO deep-learning workflow. 256x256 tiles were extracted from H&E-stained PDO while slide images (WSIs) and input into a pre-trained Hover-Net ResNet50 backbone for nuclear segmentation. In the initial transfer learning phase (Phase 0), classification weights were re-trained while backbone features were frozen. In the fine-tuning phase (Phase 1), residual layers of the ResNet50 backbone were unfrozen, and the entire network was optimized using a reduced learning rate. Model performance on segmentation was evaluated using dice coefficients across epochs, with the best checkpoint selected at epoch 32. Final classifications on tumor/normal were made at the single-cell level based on nuclear features, and PDO-level predictions were aggregated using a winner-take-all (WTA; also known as majority voting) strategy. (**C**) Representative examples of classified tumor PDOs, normal PDOs, and mixed tumor/normal PDO cultures. Individual nucleated cells are classified as tumor (red) or normal (green). (**D**) Performance of TransferNet-PDO in classifying individual cells as tumor or normal. (**E**) Performance of PDO-level classification via WTA among classified cells. For **D** and **E**: (**left**) sensitivity, specificity, and ROC-AUC across training, validation, and three independent WSI validation cohorts; (**right**) corresponding ROC curves for each cohort. Detailed methodology is described in **Supplementary Methods**.

## Methods and Materials

The study protocol was approved by The University of Pittsburgh Institutional Review Board (IRB) (IRB99-069). Participants gave informed consent to participate in the study before taking part. All samples have written informed patient consent.

A full description of materials and methods is provided in the **Supplementary Methods**. In brief, fresh tumor and matched normal tissues were collected via the institutional Head and Neck Cancer SPORE. Tissues were processed to develop PDOs, cultured in Matrigel, monitored by bright-field microscopy, and evaluated histologically. We built TransferNet-PDO leveraging Hover-Net^25^ architecture with ResNet50 backbone to distinguish tumor from normal PDOs on H&E images. Whole-exome sequencing (WES) was performed to characterize genomic alterations and clonality in tumor and PDOs, with somatic mutations identified by GATK4-MuTect2^36^ (v4.6.1.0), somatic copy number variations (CNVs) by GATK4-somatic CNV, and clonal architecture by RETCHER^37^ (accessed Jan 2025). Drug responses to cisplatin, cetuximab, lenvatinib, radiation, and chemoradiation were assessed using the CellTiter-Glo^®^ luminescent assay, which quantifies intracellular ATP as a proxy for metabolically active (viable) cells.

## Results

### HNSCC PDO models recapitulate the morphological and pathological features of the original tumor

To establish PDOs from fresh HNSCC tissues, tumor specimens were collected during surgery and transported at 4°C to our research lab (**Fig. S1A**). Time from collection to processing was under four hours for all samples, with most processed within two hours. Tumor tissues were minced, digested with trypsin, and embedded in 3D Matrigel for culture (**Fig. S1B**). Organoids retained epithelial morphology over serial passages and displayed malignant histopathologic features resembling the original tumor per pathology evaluation. Brightfield imaging showed a typical growth trajectory, from dispersed cells on day 0 to well-formed organoid spheres by day 14 (**Fig. S1C**). H&E staining and immunohistochemistry (IHC) with HNSCC morphology markers (PanCK, p63, CK13) and cell proliferation marker Ki67 (**Table S1**) confirmed squamous phenotype and histologic aggression (**Fig. S1D**). While initial cultures may contain a mixture of epithelial, immune and stromal cells, PDOs progressively became almost all epithelial, consistent with prior reports^38^.

Characteristics of the patient cohort is described in **Table 1**. After excluding one necrotic sample, tumors from 23 patients had sufficient viable cells to pursue PDO development (**Fig. S1E**). Organoid cultures were successfully established in 19 of 23 cases (82%), yielding predominantly tumor PDOs (n=14), and normal epithelial organoids overtook the culture in a minority of cases (n=5). PDOs were derived from a diverse set of tumors across anatomical sites (oral cavity, larynx, etc.), T-category (T2/3/4), and sample types (primary, recurrent, second primary) from patients of different genders, racial backgrounds, and ages (**Table 1; Fig. S1E-G**). PDOs were established from tissue inputs as small as ∼100mg (median: 366mg; **Fig. S1H**). Tumors of all 19 cases were subsequently found to be p53-mutant by IHC staining (**Table 1**). Notably, p53 status was not used as a selection criterion; all cases with sufficient viable tissue were included for this study, and p53 IHC was performed retrospectively after specimen collection. This observation reflects the high prevalence of *TP53* mutations in HPV-negative HNSCC^39^.

**Table 1.** Clinical and demographic characteristics of HNSCC patients.

| Study ID | PID | Age in years | Tumor Location | T-category (path) | Type of Tumor | Grade | Tumor P53 IHC status | Organoids status | Nutlin selection | Organoid max passage no. banked |
| --- | --- | --- | --- | --- | --- | --- | --- | --- | --- | --- |
| HN23-10728 | P001 | 47 | Larynx | T3 | Primary | POOR | Unknown | Failed | NA[1] | NA[1] |
| HN23-10730 | P002 | 75 | Larynx | T3 | Primary | POOR | Positive | Tumor | No | 12 |
| HN23-10750 | P003 | 75 | Oral cavity (floor of mouth) | T3 | Primary | POOR | Unknown | Failed | NA[1] | NA[1] |
| HN23-10752 | P004 | 58 | Larynx | T4A | Primary | MOD | Positive | Normal | No | 9[2] |
| HN24-10791 | P005 | 62 | Larynx | T4 | Unknown | MOD | Positive | Normal | No | 5[2] |
| HN24-10798 | P006 | 76 | Tongue | T2 | Unknown | WELL | Unknown | Failed | NA[1] | NA[1] |
| HN24-10810 | P007 | 54 | Oral cavity (floor of mouth) | T2 | Primary | MOD | Negative | Tumor | No | 19[4] |
| HN24-10831 | P008 | 64 | Larynx | T3 | Primary | MOD | Positive | Normal | No | 6[2] |
| HN24-10856 | P009 | 81 | Oral cavity (gum) | TX | Recurrence | MOD | Positive | Tumor | No | 8 |
| HN24-10900 | P010 | 52 | Oropharynx | T3 | Recurrence | Unknown | Positive | Tumor | No | 6 |
| HN24-10909 | P011 | 58 | Tongue | T3 | Primary | MOD | Positive | Tumor | No | 12 |
| HN24-10936 | P012 | 82 | Oral cavity (cheek) | T2 | Primary | MOD | Unknown | Tumor | Both[6] | 6 |
| HN24-10942 | P013 | 54 | Larynx | NE[5] | Primary | MOD | Positive | Normal | No | 3[2] |
| HN24-10961 | P014 | 59 | Larynx | T4a | Primary | MOD | Positive | Tumor | Yes | 7 |
| HN24-10966 | P015 | 44 | Tongue | T3 | Primary | WELL | Negative | Tumor | Both[6] | 16[4] |
| HN24-10970 | P016 | 78 | Larynx | T4a | Primary | POOR | Unknown | Failed | NA[1] | NA[1] |
| HN24-10981 | P017 | 68 | Larynx | T4a | Recurrence | MOD | Unknown | Dead tissue | NA[1] | NA[1] |
| HN24-10988 | P018 | 57 | Larynx | T3 | Primary | MOD | Negative | Normal | No | 2[2] |
| HN24-11012 | P019 | 64 | Oral cavity | T4a | Second primary | MOD | Negative | Tumor | No | 6 |
| HN24-11042 | P020 | 62 | Larynx | T4a | Second primary | MOD | Positive | Tumor | No | 2[3] |
| HN24-11074 | P021 | 64 | Oral cavity (buccal mucosa) | T4a | Primary | MOD | Positive | Tumor | Both[6] | 3[4] |
| HN25-11105 | P022 | 56 | Oral cavity (buccal mucosa) | T4A | Primary | POOR | Negative | Tumor | No | 4[3] |
| HN25-11113 | P023 | 68 | Oral cavity (floor of mouth) | T4a | Primary | MOD | Unknown | Tumor | Yes | 5 |
| HN25-11122 | P024 | 75 | Tongue | T4a | Primary | MOD | Positive | Tumor | Both[6] | 9[4] |

[1] NA = Not Applicable.
[2] Banked after normal PDOs outgrew normal PDOs.
[3] Banked early due to growth rate was slow.
[4] Active in culture. Passage number increasing at time of manuscript submission.
[5] NE = Not Evaluable. Biopsy only, therefore no path stage was evaluated per AJCC guidelines.
[6] Both = both selected and unselected cultures banked.

A frequent challenge in PDO development is overgrowth of normal epithelial cells^40^, as observed in other tumor types such as lung^38 41^ and prostate^42^ cancers. Tumor tissues frequently contain a mixture of tumor and normal epithelial cells, and in a subset of cases, these normal cells can grow rapidly and overtake tumor cells in culture. To address this issue, we implemented Nutlin-3^43^, an MDM2 inhibitor that activates the p53 pathway and selectively induces apoptosis in cells with wild-type p53 (typically normal epithelial cells) (**Fig. S2A-B**). Adding Nutlin-3 to organoid media has previously been used in many types of tumor organoids, including lung^38 44^ and liver^45^. In cases with mixed populations, this strategy achieved close to 100% malignant cell content post-selection (**Fig. S2C**). Previous studies have reported that normal epithelial cells cease proliferation after multiple passages in other tumor types^46^. In contrast, we observed that in HNSCC cultures, normal epithelial cells overgrew tumor cells as early as passage 1 (p1), and persisted throughout the rest of passages (**Fig. S2D**). In 6 of 14 tumor PDO cases (43%), Nutlin-3 was selectively applied at early passages where organoids exhibited normal-like morphology by brightfield imaging (**Fig. S2E-F; Table 1**) and subsequently confirmed by WES on lack of driver somatic mutations. These cases were more frequently associated with larger tissue specimens, which likely included a greater proportion of normal epithelial cells (**Fig. S2F**). Together, these results establish a robust and reproducible platform for generating high-fidelity HNSCC PDOs from fresh surgical specimens, setting the foundation for further analyses.

### AI model predicts malignancy in PDO cultures consistent with the original tumor

While we established PDOs from a majority of HNSCC samples, distinguishing tumor from normal or normal-like epithelial organoids remains critical for accurate interpretation of drug sensitivity experiments. Considering the relative growth of tumor *versus* normal PDOs varies across patients and specimens, this is particularly important in early passages where mixed populations are observed. As an initial benchmark, we examined the distribution of organoid size and shape. Normal PDOs appeared larger on average than tumor PDOs (9,650±299 *versus* 6,657±250 µm, maximum diameter; mean ± S.E.M., n=17 images), the difference was not statistically significant (*P*=0.11, linear mixed-effects models [LMM]; **Fig. S3A-B**). Normal PDOs typically displayed a well-differentiated cystic structure with smooth boundaries, while tumor PDOs formed irregular, solid spheres on brightfield imaging (**Fig. S2E**) and displayed abnormal nuclear features on H&E (**Fig. S3C**).

To enable classification of PDO identity at scale, we developed TransferNet-PDO, a new framework based on Hover-Net^25^ architecture and transfer learning, to classify individual cells from H&E images as tumor or normal, and assign PDO-level predictions by winner-take-all (WTA, also known as, majority voting; **Fig. 1B**). To train the model, we generated 256 x 256 tiles of high-density tumor or normal organoids (hereafter referred to as PDO large-tile datasets) from early passages, segmented nucleated cells using a pre-trained ResNet50 CNN model^25^, and manually annotated nuclei to build a balanced training dataset (1,490 tumor nuclei from 88 tiles + 1,545 normal nuclei from 78 tiles). After feature transfer (phase 0) and full model fine-tuning (phase 1), we selected the best-performing model on the classification task by ROC-AUC (area under the receiver-operating characteristic curve) (**Fig. S4A; Fig. 1B**; examples shown in **Fig. 1C**).

After training, the model was validated using three distinct cohorts of whole slide images (WSIs; hereafter referred to as PDO WSI datasets), each designed to evaluate a different aspect of model generalizability (**Fig. S4B**). Specifically, we used: (1) non-overlapping regions (∼87.5%) from the same WSIs where PDO large-tiles were selected, to assess the model’s ability to extend to unseen areas within the same images (34,859 cells); (2) WSIs from later passages of the same specimens, to evaluate performance on temporal variation in PDO morphology (25,625 cells); and (3) WSIs from entirely different patients, to test model generalization to unseen individuals (19,570 cells) (**Fig. S4B**; **Fig. S5**). At the single-cell level, TransferNet-PDO achieved ROC-AUCs of 0.94, 0.81, and 0.78 on these datasets, respectively (**Fig. 1D**). At the PDO level, we applied a WTA approach, assigning each PDO’s class based on the majority cell type prediction, which yielded ROC-AUCs of 1.00 on the same images, 0.90 within the same patients, and 0.88 across different patients (**Fig. 1E**). Together, these findings demonstrate that TransferNet-PDO enables accurate, salable, and robust classification of tumor versus normal PDOs directly from H&E slides, offering a practical tool for enhancing the reliability of organoid-based research and potential clinical applications.

### Whole-exome sequencing identifies consistent genomic alterations between HNSCC PDO models and the original tumor

To assess the genomic fidelity of PDOs relative to their original tumors, we performed WES on frozen tumor specimens, matched normal tissues, and serial PDO passages (∼200X per sample after duplicate removal), to identify somatic mutations in individual tumor and PDO with normal as control. We focused on WES data from six cases where Nutlin-3 selection was not applied, where tumor clonality would only be a function of selection pressure under baseline organoid culture conditions (**Table S2**).

Tumor mutational burden (TMB) remained consistent between PDO passages and their original tumors (**Fig. 2A**, bar plot above the heatmap), with minor fluctuations due to gain or loss of low-frequency passenger mutations. Driver mutations remained stable over time, and non-overlapping variants consisted of mostly passenger mutations, consistent with literature^20^. Across all PDO passages, protein-altering somatic mutations in key driver genes associated with HPV-negative HNSCC^39^, such as *TP53, CDKN2A, MLL2, CUL3, NSD1, PIK3CA, NOTCH1, NOTCH2, EGFR*, were consistently retained with minimal disruption compared to the original tumor (**Fig. 2A**; **Table S3**; full somatic mutation lists in **Table S4**). Each patient harbored distinct *TP53* mutations (e.g., P190L, P278R, P278S, P213*, P248Q, R238W). In HN24-10909, two *TP53* mutations (P278S, P213*) co-existed. In HN24-10909 and HN24-10936, low tumor cellularity in the original sample potentially resulted in reduced detection of somatic mutations by WES; these were reliably detected in tumor-derived PDOs. *TP53* mutation status from WES were verified by p53 IHC (**Fig. 2B**, p53 positive/negative examples shown). In contrast, normal PDOs exhibited wild-type p53 expression by IHC and lacked any recurrent somatic driver mutations by WES. Once tumor cells became dominant, they rapidly expanded and persisted with high malignant cell content across passages, as reflected by *TP53* somatic variant allele fractions approaching 100% (**Fig. S6A**). One exception was HN24-10936, where malignant cell populations increased in PDO over time yet remained mixed with normal epithelial cells. Overall, these molecular findings aligned with our pathological observations, suggesting that tumor *versus* normal identity was often established early in culture.

**Figure 2.**
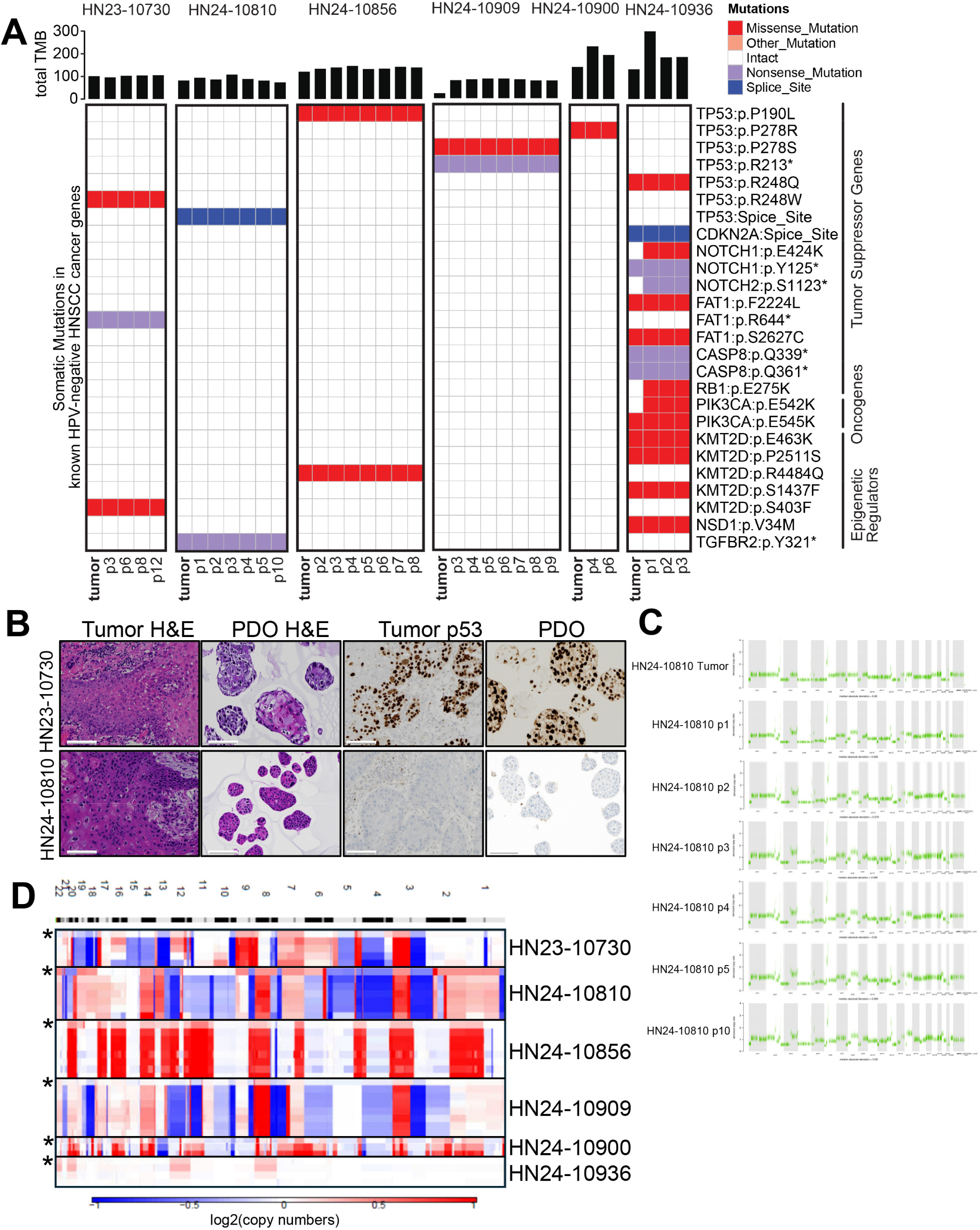
HNSCC PDOs show high-fidelity genomic landscape. (**A**) Somatic mutations in known HNSCC cancer genes across six HPV-negative HNSCC patient tumors and corresponding PDOs across serial passages (indicated as p#). (**top**) bar plots show total tumor mutational burden (TMB) per sample, defined as the number of protein-altering somatic variants (SNVs and small insertions/deletions [indels]) by whole-exome sequencing (WES); (**bottom**) the heatmap displays mutations in known genes involved in HNSCC cancer development and progression, categorized by functional class: tumor suppressors, oncogenes, epigenetic regulators, and others. Each row indicates a unique variant shown to the right of the heatmap; each column represents a tumor or PDO sample. Variant classifications are color-coded. (**B**) Representative H&E and p53 IHC staining of tumor tissues and matched PDOs from two patients (HN23-10730 with positive p53, and HN24-10810 with negative p53). PDOs show histologic and immunohistochemical concordance with the original tumor. (**C**) Representative example of tumor and serial PDO passages from patient HN24-10810 (**top to bottom**: p1-p10) showing genome-wide somatic copy number variations (CNV). Denoised copy ratio is shown on *y*-axis. Chromosome numbers are shown on *x*-axis. CNV patterns remain largely stable across passages, reflecting genomic fidelity. (**D**) Heatmap of log_2_ copy number changes across the genome for all six patient-PDO pairs from **A**. Each row represents a tumor (denoted by asterisk, first row of each panel) or PDO sample; chromosome number and cytobands are shown on the *x*-axis. Gains (red) and losses (blue) are shown relative to diploid baseline, demonstrating preservation of major CNV events in PDOs.

Somatic CNV analysis further supported the genomic stability of PDOs, showing strong concordance between the original tumors and PDO passages (**Fig. 2C-D**; **Table S5**). While WES data were analyzed for PDOs within 12 passages, H&E images of long-term passages in p14 showed similar pathology to earlier passages (**Fig. S6B**). Taken together, these findings demonstrate that HNSCC PDOs faithfully preserve the mutational landscape of the original tumor across serial passages, reinforcing their genomic stability and translational relevance for tumor modeling, drug testing, and precision oncology applications.

### HNSCC PDO models preserve clonality architecture across serial passages

Upon confirming the consistent p53 mutant status from early to late passages (**Fig.3A**), we sought to evaluate whether HNSCC PDOs preserve the global clonal architecture of their original tumors. Retention of subclonal diversity is critical for modeling intratumoral heterogeneity and for using PDOs as preclinical tools to study therapeutic response. To address this question, we reconstructed subclonal structures using integrated somatic mutation and CNV data (**Fig. 3B**;**Table S6**). In five out of six sequenced cases, the founding clone was *TP53*-driven and remained consistently across PDO passages (**Fig. 3B**). In one exception (HN24-10900, oropharynx), *TP53* was detected as a subclone in the original tumor. Organoids of this case required antifungal treatment to eliminate a fungal infection from tumor (**Fig. 3B**, blue arrow), which altered the clonal dynamics and led to the loss of a minor subclone (**Fig. 3B**, yellow node on the tree). Despite this, the *TP53* subclone persisted and drove organoid growth, preserving malignancy in culture.

**Figure 3.**
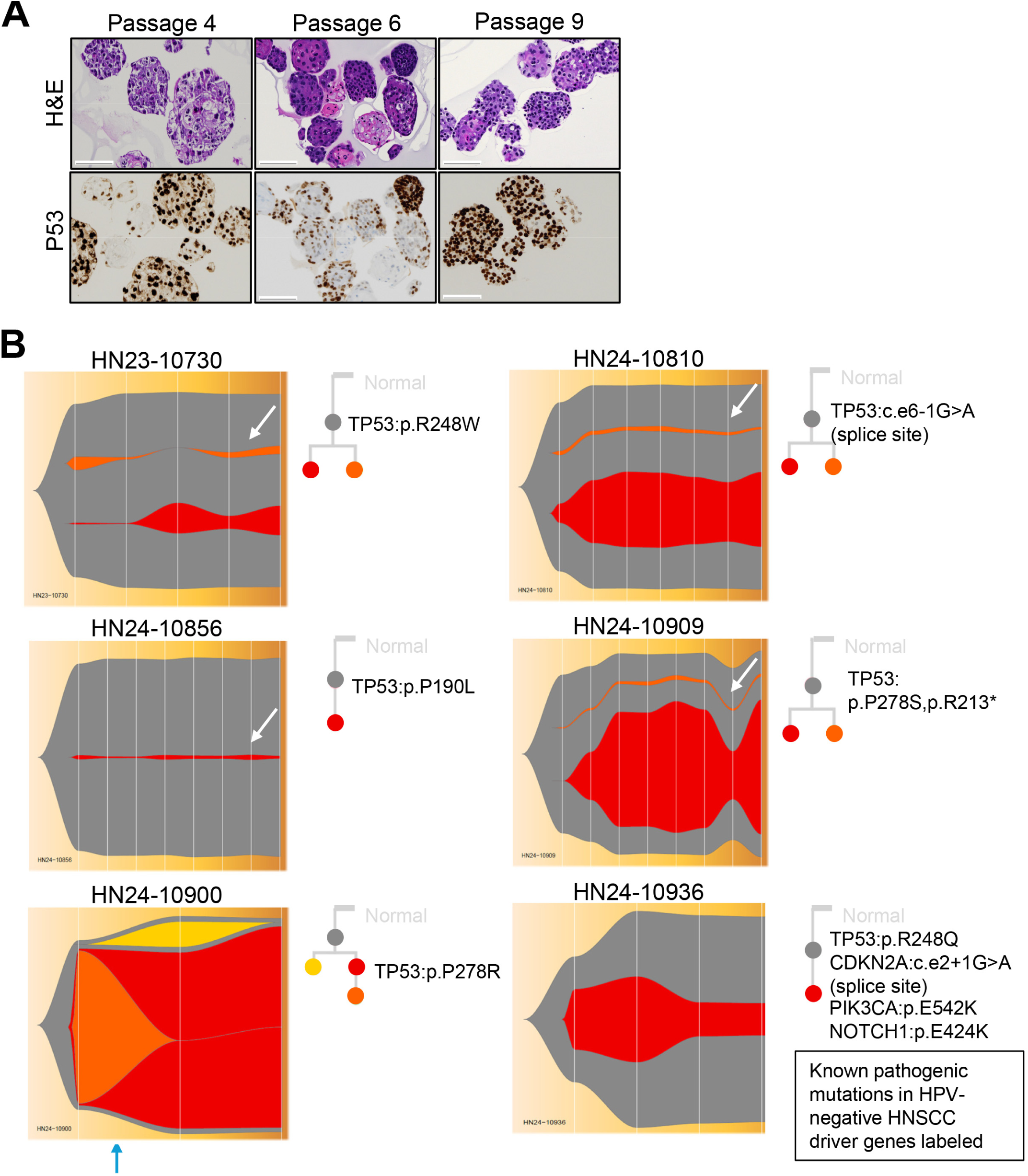
HNSCC PDOs maintain clonal architecture over time. (**A**) Representative example of H&E and p53 IHC staining of PDOs from patient HN23-10730 across passages 4, 6, and 9. PDOs retain malignant histology and consistent nuclear p53 accumulation over time, reflecting the underlying *TP53* mutational status. Scale bar = 100 μm. (**B**) Clonal architecture reconstruction from WES somatic mutation and CNV analysis for six HPV-negative HNSCC cases from Fig. 2. Fishplots depict subclonal dynamics from the original tumor across PDO passages, with each vertical line representing a sample in the same order from Fig. 2A. *TP53* mutant clones were the founding clones in 5 of 6 cases. Case HN24-10900 showed a clonal shift after antifungal treatment (blue arrow). Minor subclones (white arrows) were preserved. Known pathogenic mutations in HNSCC driver genes are annotated next to each plot, with colored dots on the phylogenetic tree corresponding to specific subclones from the fishplots.

Among the other five cases, we observed subclonal evolution within the dominant *TP53* clone. For example, in HN24-10936, a *PIK3CA/NOTCH1*-driven subclone emerged and remained stable across passages. Subclones of low prevalence (<5% abundance) were retained in multiple cases (**Fig. 3B**, white arrows), suggesting that minor subpopulations from the original tumor can be retained in serial PDO cultures.

### HNSCC PDO models demonstrate sensitivity to chemotherapy, radiation, chemoradiation, and targeted therapy

Building on their preserved histologic and genomic fidelity, we next evaluated the drug sensitivity profiles of HNSCC PDOs. Stable tumor organoid cultures were treated with cisplatin (chemotherapy), cetuximab (anti-EGFR), lenvatinib (a multiple RTK inhibitor against VEGFR1-3) (**Table S7**), radiation, and chemoradiation, and measured for cell viability. Cisplatin treatment induced a dose-dependent reduction in cell viability as measured by the CellTiter-Glo assay, with IC50 values ranging from 1.61 to 15.69 µM across cases (**Fig. 4A**; IC50 listed in **Table S8**). In contrast, cetuximab had a minimal effect on PDO viability (**Fig. 4B**). This finding is consistent with literature reporting response to cetuximab is heterogeneous and often limited in HNSCC^47^. Similarly, lenvatinib did not significantly reduce cell viability across a range of doses, except at very high concentrations, likely due to off-target toxicity (**Fig. 4C**).

**Figure 4.**
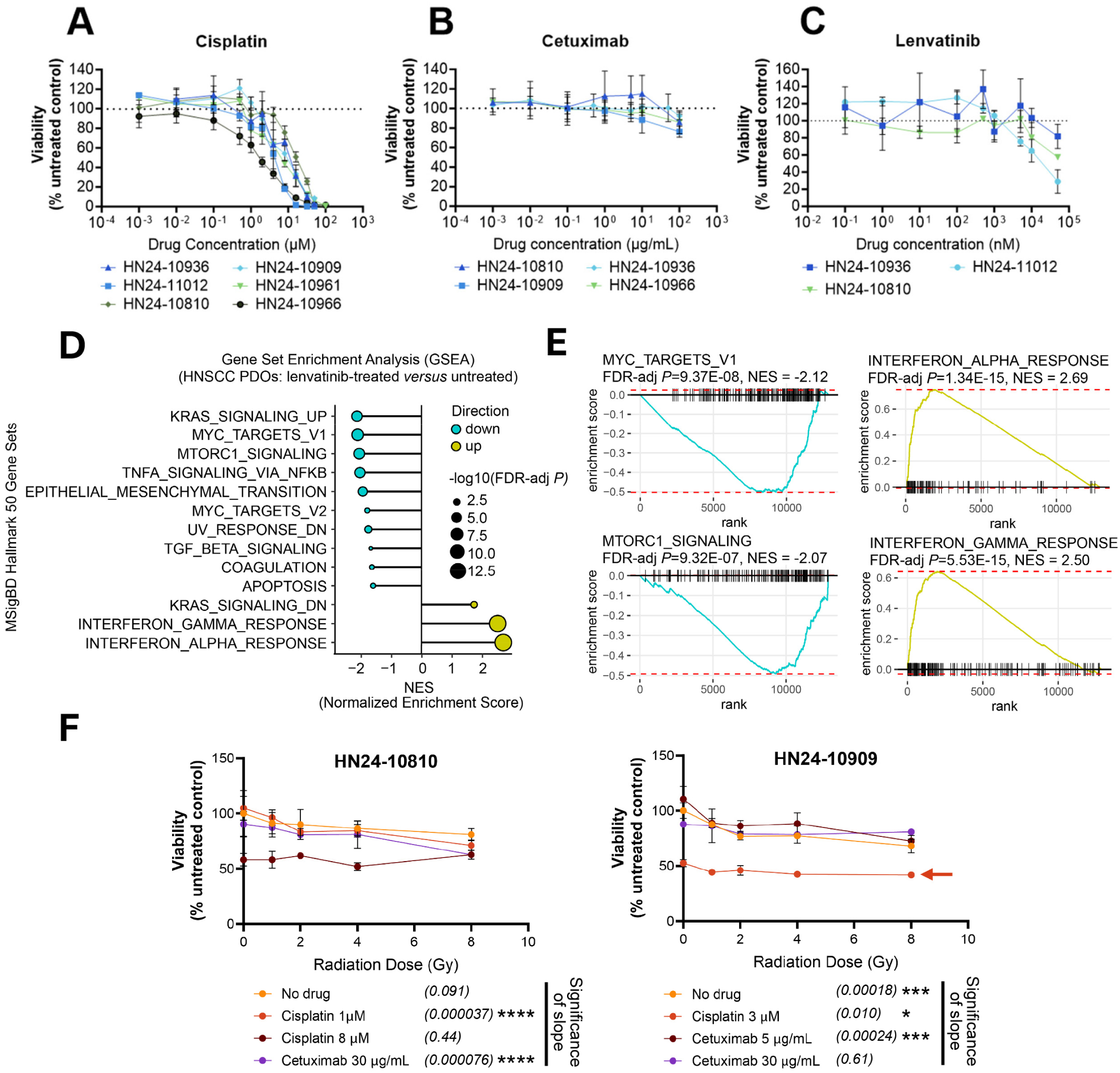
Tumor PDOs demonstrate clinically relevant sensitivity to standard-of-care therapies in HNSCC. (**A-C**) Dose-response curves for cisplatin in **A**, cetuximab in **B**, and lenvatinib in **C** on PDOs from multiple patients. PDOs showed consistent sensitivity to cisplatin (IC50 range: 1-20 μM), while cetuximab and lenvatinib had minimal effects at most tested doses. Viability was measured using the CellTiter-Glo^®^ assay and normalized to untreated control. (**D**) Gene set enrichment analysis (GSEA) of RNAseq gene expression from Lenvatinib (10^2^ nM, 24 hours)-treated PDOs (FDR-adjusted *P*<0.05) from four patients (eight samples total). Top MSigDB Hallmark 50 gene sets (H50) are shown on the row. Dots are colored by direction (NES>0: upregulated; NES<0: downregulated), and their size are scaled by -log10(FDR-adjusted p-value). A full list of H50 GSEA results is provided in **Table S9**. (**E**) GSEA plots showing lenvatinib induced downregulation of MYC targets and OXPHOS, and upregulation of type I/II IFN signaling in HNSCC PDOs. Enrichment scores are shown on *y*-axis, and ranked gene positions are shown on *x*-axis. (**F**) Chemoradiation response in two tumor PDO lines. For each treatment group, four replicates were included. P-values in parathesis next to each treatment group represent whether the slope of fitted models (indicating radiation dose-dependent cell viability reduction) was significantly different from zero within that group (full statistics are provided in **Table S10**). Denotation: * *P*<0.01, *** *P*<0.001, **** *P*<0.0001. Two-sided hypergeometric test was used in **D**. Linear mixed-effects models were used in **F**, with radiation dose as the fixed effect and replicate id as the random effect. P-values shown in **D** and **F** were after multiple comparison adjustments by BH-FDR method.

Lenvatinib targets angiogenic signaling via VEGF pathway blockade and is being investigated in combination with immunotherapy in ongoing clinical trials. To better understand its biological impact in PDOs, we performed RNAseq on treated and untreated samples from four PDO lines, followed by gene set enrichment analysis (GESA) (**Table S9**). Lenvatinib treatment significantly downregulated proliferative and metabolic programs, including MYC targets, mTORC1 signaling, among others (FDR-adjusted *P*<0.05; **Fig. 4D**). In contrast, inflammatory and immune-related gene sets, such as interferon alpha and gamma response signatures, were upregulated (**Fig. 4E**). These findings are consistent with prior reports of RTK inhibitor mediating both anti-proliferative and immune-modulating effects in human tumors^48^.

Given that chemoradiation is a standard treatment for locally advanced or recurrent HNSCC, we examined its effect on tumor PDOs to assess their utility as a translational model (**Table S10**). In HN24-10810 PDO line (**Fig. 4F, left**), radiation alone (no drug) reduced cell viability (FDR-adjusted *P*=0.091, by LMMs). Combining radiation with low-dose cisplatin (1 µM) or cetuximab (30 µg/mL) yielded a significant reduction in viability (*P*<0.0001), whereas this was not observed with high-dose cisplatin (8µM; *P*=0.44). In HN24-10909 (**Fig. 4F, right**), radiation alone significantly reduced viability (*P*=0.00018). Cisplatin alone (0 Gy) reduced viability, and increasing radiation dose to 8 Gy further decreased viability by 5-10% (*P*=0.010; **red arrow**), indicating an additive chemoradiation effect. A similar radiation dose-dependent viability decline occurred with low-dose cetuximab (5 µg/mL) but not with high-dose cetuximab (30 µg/mL). Patient HN24-10909 received adjuvant chemoradiation following surgery. However, due to the timing of therapy, we could not isolate the clinical benefit of chemoradiation from surgical outcomes. Collectively, these findings show that HNSCC PDOs respond to radiation and chemotherapeutic agents in a treatment- and dose-dependent manner, supporting their potential as a platform for functional testing of therapeutic regimens.

## Discussion

We established PDOs from a diverse cohort of HPV-negative HNSCC patients, representing various anatomical sites, tumor T-categories, and demographics. We created TransferNet-PDO, a novel transfer-learning based CNN on rapid and accurate classification of organoid identity (malignant *versus* normal) at single-cell resolution, archiving high performance across passages and patients (AUC≥0.88). This AI tool enables morphological tracking over time and with treatment, which can be adapted to support dynamic drug response monitoring, functional screening, and multi-center biobanking. Future iterations incorporating multimodal data^49^ (e.g., transcriptomics, proteomics) and histology foundation models^29–34^ with fine tuning may further improve classification and predict therapeutic prediction.

Our platform preserves genomic stability and clonal architecture across serial passages, supporting PDOs as reliable models for personalized therapy. While prior studies have reported genomic shifts in culture^18^, our focus on short-term passages (<12; 3 months) preserves key genomic features, with ongoing monitoring to assess long-term stability. Short- and long-term PDOs offer complementary strengths for translational research. Short-term PDOs offer a practical platform for real-time treatment outcome prediction and monitoring as patient “avatars”, while long-term cultures enable modeling of tumor evolution and resistance through large-scale drug screening.

Drug sensitivity assessment demonstrates the translational relevance of our PDO models. Cisplatin reduced PDO viability in a dose-dependent manner, while cetuximab and lenvatinib did not eliminate tumor organoids, aligning with known clinical scenarios. As an NAM platform, PDOs offer the potential to model therapy resistance, such as clonal selection in the TME, which may advance our understanding of treatment failure. Incorporating PDOs from longitudinal tumors upon recurrence or progression may further strengthen the impact of this model in interrogating preexisting or acquirement of resistant clones.

While various PDO technologies exist, the constitutive PDO system offers distinct advantages for genetic and pharmacologic manipulation, enabling precise control over specific cell types^50^. HNSCC PDOs can be engineered to express specific alterations (e.g. NOTCH1/2 mutations^51^) to study drug-resistant subclones, with the potential to guide rational combination therapies with novel small-molecule or immune checkpoint inhibitors^52^. Additionally, the platform provides a tractable system to evaluate CAR T cell efficacy^53 54^ in a personalized context, including antigen targeting, cytotoxicity, and mechanisms of immune evasion.

Our study has limitations. The patient cohort, while diverse, sample size may limit generalizability across the broader HNSCC population. For patients who received surgery without further treatment, direct correlations between PDO responses and clinical outcomes require long-term monitoring upon progression. Nutlin-3 selection, selectively applied when normal cell overgrowth occurred, is limited to p53-mutant tumors^14^. Prior studies also suggested trypsin may affect long-term genomic stability of PDOs^18^, and alternative culture conditions, such as TME-mimicking extracellular matrix scaffolds^55^ or hypoxia^56^, could mitigate this risk, with orthogonal validation (e.g., single-cell RNAseq) to confirm clonal stability. Our AI classifier, while robust, relies primarily on nuclear and regional features and may underutilize cytoplasmic information. 3D imaging or single organoid microengineered devices^53^, may enable real-time, label-free tracking of growth and drug response.

This work opens several translational avenues. Longitudinal PDOs from sequential samples could track tumor evolution and resistance, supporting real-time adaptation of therapeutic strategies, for instance, timely transition from immunotherapy to chemotherapy or when chemotherapy should be added prior to clinical progression in recurrent HNSCC. Furthermore, assembloids incorporating tumor, immune, and stromal components may better model the TME and enable identification of patient-specific immunotherapy biomarkers or functional validation of computationally predicted drug combinations. Pairing these models with AI tools trained on multi-omic PDO data will offer a path forward toward scalable, high-throughput therapeutic prioritization. Lastly, our finding that PDOs can be developed from as little as 100mg of tissue marks the feasibility for biopsies, particularly in recurrent, inoperable cases or metastatic cases, where a better understanding of drug sensitivity and resistance evolution could be transformative.

Beyond HNSCC, our PDO platform has broader implications for precision oncology. The approach is adaptable to other heterogeneous solid tumors, such as pancreatic or lung cancers, where predictive biomarkers remain elusive. Integrating PDOs with AI-driven analytics provides a generalizable framework for drug repurposing, combination therapy design, and biomarker discovery across cancer types on rapid translation of personalized therapies from bench to bedside.

## Conclusion

We established a clinically relevant PDO platform for HPV-negative HNSCC that preserves tumor fidelity, enables drug response modeling, and supports AI-driven classification. These integrated tools demonstrate the utility of PDOs for translational research and provide a foundation for developing complex assembloids and personalized therapeutic strategies in head and neck cancer.

## Author Contributions

R.B. conceptualized the study. R.B. and J.J.L. designed the study. R.D. developed and performed the experiments, selected and optimized the protocols, and built the PDO platform. J.C. and B.I. developed and validated the AI models. C.K. and K.B.S. assisted with PDO development. J.D.S. contributed surgical samples, provided surgical insights, and coordinated on the surgical side for specimen workflow. M.H. performed chart reviews of patients. B.E.D. assisted with experimental design. D.D.B shared early versions of PDO protocols and was critical in providing feedback and sharing experience throughout this study. A.V.L. contributed resources and guidance in PDO development. X.W. contributed AI insights. M.E.S., S.K., S.S., K.C. consented patients and contributed surgical specimens. K.S. coordinated specimen processing for H&E and IHC experiments and performed slide scanning. C.R. assisted with clinical annotation of patients. M.L. coordinated and performed WES QC and library preparation. J.L. and S.L. coordinated and performed RNAseq QC and library preparation and sequencing. D.P.Z. contributed to clinical translation discussion and provided critical insights into HNSCC trials. H.D.S assisted with coordination of specimen collection workflow via HN SPOPE. I.K.Z provided critical insights into ex vivo experiments. L.V. contributed to study design, protocol optimization, and data interpretation. R.L.F. and J.P.Z. contributed samples and resources, and provided support on ensuring stability of surgical specimen workflow. R.R.S led biospecimen collection via HN SPORE, performed pathology reviews, and provided critical insights from pathology evaluation. R.D., J.J.L., and R.B. wrote the manuscript. J.J.L. and R.B. edited the manuscript. All authors reviewed and contributed to the final manuscript.

## Acknowledgement

We thank the patients and families for their participation in this study. We thank Dr. Fangping Mu for technical assistance at The University of Pittsburgh Center for Research Computing (CRC) high-performance computing clusters (HPC). We thank Dr. Laura Andres-Martin (The New York Stem Cell Foundation Research Institute) for generous sharing of resources in early development of the project. This work was supported by National Institutes of Health (NIH) grant R01DE031729 (R.B., J.J.L.), head and neck cancer SPORE P50CA097190 (R.B., R.L.F., J.P.Z., D.P.Z., H.D.S.), UM1CA186690 (J.J.L.), Cancer Immunology Training Program T32CA082084 (R.E.D.), Postdoctoral Training in Head and Neck Oncology T32CA060397 (J.D.S., J.P.Z.), and in part by National Cancer Institute (NCI) through the UPMC Hillman Cancer Center (HCC) CCSG award P30CA047904 (R.L.F., R.B.), and The University of Pittsburgh CRC through the resources provided, specifically the HTC clusters supported by NIH S10OD028483. This project used the UPMC HCC Cancer Bioinformatics Facility (CBS), Cancer Genomics Facility (CBF), and Translational Oncologic Pathology Services (TOPS). **<u>Role of funding sources</u>**. The funding sources had no role in the study design, data collection, data analysis, interpretation, or writing of the manuscript.

