## Supplementary material for "Integrating Artificial Intelligence-Driven Digital Pathology and Genomics to Establish Patient-Derived Organoids as a Novel Alternative Model for Drug Response in Head and Neck Cancer": Supp Methods and Supp Figs 1-6

This document contains the full description of **Materials and Methods** for the manuscript, **Supplementary Figures 1 to 6, Title of Supplementary Tables 1-10**. **Content of Supplementary Tables 1-10** is provided in a separate spreadsheet, and also available at GitHub repository (link: https://github.com/HCC-data-sciences-pub/HNSCC-PDO-integrative-analysis).

### Supplementary Materials and Methods

#### Human specimens

The study protocol was approved by The University of Pittsburgh Institutional Review Board (IRB)-approved protocol (Protocol No. 99-069). Participants gave informed consent to participate in the study before taking part. All samples have written informed patient consent. Tumor and normal tissue were collected during surgical resection or biopsy via the Head and Neck Cancer SPORE at the University of Pittsburgh.

#### Media

The base media used was advanced DMEM/ F12 (Gibco) supplemented with 1% Penstrep, 1% HEPES, and 1% GlutaMAX (abbreviated as adDMEM/F12+++). Other media components were added to produce two organoid growth media recipes, termed HN media[1] and M7 media[2].

#### Head and neck tumor tissue processing

The workflow is illustrated in **Fig. S1**. Tissue specimens were placed into chilled adDMEM/F12+++ medium. At the time of tissue collection, peripheral blood was collected into heparinized tubes and PBMCs were isolated from the blood. For the tumor and normal tissue, sections of tissue were flash frozen for whole exome sequencing, and additional sections were fixed in formalin for histochemical analysis. The remaining tumor tissue was processed according to the method previously described by Driehius et al[3 4]. Briefly, tissue was minced into small pieces with a sterile scalpel and digested in a solution of 0.125% trypsin in adDMEM/F12+++ medium at 37°C to produce a suspension of cell clusters. Digestion was monitored closely, and the tissue suspension was mixed by pipetting every 10 minutes. Digestion was ended either when the tissue was fully broken down into small clusters of cells or after 60 minutes, whichever came first. At that time, the digest suspension was poured over a 100 µm strainer, and cells and small clusters were collected by centrifugation at 500 x *g* for 5 minutes. This cell pellet was washed once with adDMEM/F12+++ medium, then the cell pellet was re-suspended into 70% Cultrex Basement Membrane Extract (BME) (R&D Systems). Cells in 10 µL drops of BME were plated into 24-well suspension culture plates, and after allowing the gel to solidify for 15-30 minutes at 37°C, 500µL of organoid medium (either HN or M7 medium) was added per well. The plates were then transferred to the cell culture incubator at 37°C.

#### Head and neck organoid culture

After plating cells from the tumor, the organoid medium was refreshed every 2-3 days, and organoids were passaged with TrypLE approximately every two weeks. Organoid growth was monitored by brightfield microscopy. After organoids were grown either to confluence or when organoid diameter exceeded 200 μm, organoids were passaged by digestion in TrypLE (Gibco), then plated in fresh BME drops. Passage ratios varied from 1:3 to 1:8 depending on density of organoids. Initially, half of the wells were cultured with HN medium and half were cultured with M7 medium, and medium was kept consistent throughout serial passages per PDO line, which was important for WES analysis. Details of media compositions were described as cited: adDMEM/F12+++ media recipe[1], HN media recipe[1], and M7 media recipe[2]. Our observations were consistent with previous work showing that while organoids grew faster in M7 medium compared to HN medium[2], the media used did not affect organoid morphology or genomics[5]. At the time of passage, a fraction of the single cell suspension was viably frozen or snap frozen. Every 2-3 passages, a fraction of organoids was fixed and embedded in paraffin for histochemical analysis and imaging.

**Hematoxylin and eosin (H&E), immunohistochemistry (IHC) staining, and image scanning**

On the day of receiving tissue samples, tumor and normal tissue were fixed in 10% neutral buffered formalin overnight, then dehydrated and embedded into paraffin blocks at the UPMC Clinical Test Development Laboratory. For organoid samples, Matrigel was removed using organoid harvesting solution (R&D Systems) according to the manufacturer's protocol, then organoids were fixed in 10% NBF and pelleted in a 1.5mL conical tube. Histogel was then added to the organoid pellets, which were embedded into paraffin blocks. Slide thickness was 4 microns. For p53 and Ki-67 staining, immunohistochemical staining was performed using the Leica Bond III autostainer at the pathology department at UPMC. For all other antigens, immunohistochemical staining was performed manually (**Table S1**). Slides were imaged at 10X magnification on the Nikon NI-E brightfield microscope. H&E and IHC images were scanned at 40X by Leica AT2 scanner. p53 immunostaining was used as a surrogate marker for *TP53* mutation status, recognizing that aberrant expression patterns (either diffusely strong or complete absence) may suggest underlying mutations.

**Nutlin-3 selection in PDO cultures**

The goal of Nutlin selection was to eliminate normal cell contamination from the culture. Since normal cells overgrow tumor cells within the first few passages in some cases, Nutlin selection was performed during passage 1. Nutlin-3 (Cayman Chemical, cat# 18585, stock concentration=10 mM in dimethyl formamide) was added to the organoid media at a final concentration of 10 μM. After three days, death of wild-type or normal organoids was visible by eye, while pure tumor organoids were unaffected. At this time, medium was refreshed and could be replaced by medium without Nutlin. Growth of organoids with and without Nutlin treatment was monitored by brightfield microscopy (**Fig. S2**). After selection, the remaining tumor organoids were passaged and expanded, resulting in a culture of pure tumor organoids. To ensure that tumor cells expressing wild type p53 were not accidentally eliminated, a subset of organoids not treated with Nutlin was maintained and biobanked.

**TransferNet-PDO training and validation datasets**

Training and validation datasets for TransferNet-PDO were derived from hematoxylin and eosin (H&E) whole slide images (WSIs) of patient-derived organoids (PDOs) scanned at 40X magnification (0.25 μm/pixel). For the training dataset, large tiles were extracted from WSIs and manually annotated by a pathologist (R.R.S.) to generate single-cell-level labels (tumor or normal) for each cell within the tiles. For the validation dataset, H&E WSIs were first evaluated by pathologists to assign preliminary tumor or normal labels to individual PDOs. These labels were further confirmed using p53 immunohistochemistry (IHC) on adjacent tissue slides, leveraging p53 overexpression (p53-positive mutant) or lack of expression (p53-negative mutant) as a marker of malignancy. In a subset of PDO cultures with near-100% tumor or normal cell purity, as determined by pathologist evaluation of H&E morphology, whole-exome sequencing (WES) was utilized to validate labels by identifying cancer-specific mutations (e.g., TP53) in tumor PDOs or their absence in normal PDOs. For mixed PDO cultures containing both tumor and normal cells, bulk WES was unreliable for distinguishing cell types; thus, labels were assigned based on pathologist's evaluation of H&E WSIs and p53 IHC at the individual PDO level as the ground truth.

**TransferNet-PDO convolutional neural network (CNN) training and validation**

We describe the development of TransferNet-PDO in three parts described as follows: 1) training dataset generation. We selected regions from 7 whole slide images (WSIs), where majority of cells inside those regions are either tumor cells or normal cells. Then, 256 x 256 tiles were generated from those regions using QuPath software (version: 0.4.4). To annotate nuclei of the generated tiles, we first segmented nuclei on those tiles using default Hover-Net ResNet50 model[6], then we manually annotated the nuclei into either normal or tumor type, with ambiguous ones excluded. With tiles and nuclei annotations, we generated modeling patches in the format required by hover-net framework. Briefly, each patch is a 5-dimensional numpy array, where the first 3 dimensions are R,G,B channels of the tile, the 4th channel is the ground truth of nucleus instances, where 0 indicates background, and non-zero integers indicate each nucleus, the 5th channel is the ground truth of nucleus pixel types, where 0 indicates background, 1 indicates normal type, 2 indicates tumor type. We randomly selected 88 tumor tiles and 78 normal tiles as training sets which totally contains 1490 tumor nuclei and 1545 normal nuclei, that left us the validation set of 22 tumor tiles (423 tumor nuclei) and 52 normal tiles (878 normal nuclei).

2) model optimization and selection. To train a new hover-net model, we utilized training API of the published hover-net framework. Briefly, training process was split into two phases. In the first phase, layers of residual blocks were frozen, and model was trained for 50 epochs, then training entered into the 2nd phase, where layers in residual blocks were unfrozen to allow updating all parameters for another 50 epochs of

training. We followed instruction of hover-net framework to set training parameters. In the config.py file, model_mode was set to 'fast', and nr_type was set to 3. We left the parameters defined in opt.py unchanged, so Adam optimization with initial learning rate of 0.0001 was used for optimization, and decay of learning rate was carried out for every 25 epochs. For batch sizes, the train and validation splits were set to 16 in the first phase, while the batch_size of train was set to 4, and validation split was set to 8 in the 2nd phase. For data augmentation, we used hover-net's transforms with modified parameters, basically, in train_loader.py, we changed the range parameter of add_to_hue from (-8,8) to (-32,8), range parameter of add_to_saturation from (-0.2,0.2) to (-1,1), range parameter of add_to_brightness from (-26,26) to (-2,2), range parameter of add_to_contrast from (0.75,1.25) to (-0.5,1.5).

3) model evaluation. After training finished, hover-net framework outputs dice scores for each epoch in both phases, we selected the best model based on those dices scores. For this particular model, we found the checkpoint at epoch 32 in phase 1 is the best, it yields dice score of 0.926893 for normal nuclei and 0.877372 for tumor nuclei from validation tiles. To further evaluate this model on the nucleus-level, we ran predictions on both training and validation tiles using the inference API of hover-net framework, with option set to 'tile', we then computed performance metrics and ROCAUC with the resulted predictions and ground truth. To evaluate our model's generalization, we further explored its performance on whole side images (WSIs). We grouped those WSIs into 3 cohorts, cohort #1 are slides where our training tiles came from, those slides contain about 12.4 % cells that were used in model training process; cohort 2 are slides from same patients of cohort 1, but are different passages of PDOs; cohort 3 are slides from different patients. We then ran prediction with our model using the inference API of hover-net framework with option set to 'wsi' to get nuclei-level predictions. For the ground truth of the nucleus types on those WSIs, we denote that all the nuclei detected by our model that resided in a particular PDO will have the same type as this PDO, for example, nuclei detected within a tumor PDO were assigned tumor type. We then generated performance metrics and ROCAUC for each cohort. For PDO-level prediction and evaluation, the predicted type of a PDO was generated by majority voting among nuclei within this PDO that were detected and classified by our model. The performance metrics and ROCAUC on PDO-level were then generated for each cohort.

**Whole Exome library preparation and sequencing**

The DNA extraction and library preparation were performed at UPMC Hillman Cancer Center Cancer Genomics Facility (GCF), and sequencing was performed at Novogene Inc, following the same procedure as we published previously[7]. DNA was extracted from frozen samples (tissues or PDOs) and subject to whole-exome sequencing (WES) library preparation using the Agilent Sure Select XT HS2 kits with human all exon V8 baits. Libraries were pooled and sent to Novogene Inc. to sequence on a NovaSeq X Plus instrument (Illumina Inc.). Paired-end (PE) 150bp reads were generated to yield on average 200X coverage for tumor, matched normal, and PDO samples.

**DNA extraction and QC.** Frozen tissue/organoids were mechanically homogenized in 360 µL of Qiagen Buffer ATL (Qiagen, Germantown, MD, catalog #: 939011), and 40 µL of QIAGEN Proteinase K (catalog #19131). DNA was extracted using the DNeasy Blood & Tissue Kit (Qiagen, Germantown, MD, catalog: 69504). Samples were incubated in a shaking heat block at 56 °C overnight. DNA extraction was performed according to the manufacturer's instructions except for using an excess volume of Buffer ATL and Proteinase K. gDNA concentration was determined using the double stranded DNA High Sensitivity Qubit kit (ThermoFisher Scientific, Pittsburgh, PA, catalog: 32854) and the gDNA quality was determined using the NanoDrop and the Agilent High sensitivity DNA Bioanalyzer kit (Agilent, Santa Clara, CA, catalog: 5067-4626).

**WES library prep protocol.** 200 ng of gDNA underwent exome library prep using the SureSelect XT HS2 DNA Library Preparation and Target Enrichment kit (Agilent, Santa Clara, CA, catalog: G9981A) assay for library construction according to the manufacturer's instructions (SureSelect XT HS2 DNA Library Preparation and Target Enrichment (Agilent, Santa Clara, CA, Version E0, July 2022). Specifically, 0.25ng/ul (50 ul low TE) ds gDNA was pipetted into Covaris microtube AFA Fiber pre-slit snap cap (6x16) tubes (Covaris, Woburn, MA, catalog: 520045). Samples were sheared using the Covaris S-1 (Covaris, Woburn, MA) with the following settings (Duty Cycle: 10%, Intensity: 5, Cycles per burst: 200, Time: 8 cycles of 40 seconds for a total of 320 seconds, Set Mode: Frequency Sweeping, Temperature: 4°C to 7°C). Achieving an average fragment size of approximately 150 base pair. The Agilent High sensitivity DNA Bioanalyzer kit (Agilent, Santa Clara, CA, catalog: 5067-4626) was used to confirm the fragment size of each sample. Once confirmed, DNA was end repaired, A's added, ligated, and underwent pre-hybridization PCR (98°C 2 minutes, twelve cycles of: 98°C 30 seconds, 65°C 30 second, 72°C 1 minute, 72°C 1 minute). The concentration and quality of the pre-hybridization PCR product was determined using the double stranded High Sensitivity Qubit kit (ThermoFisher Scientific, Pittsburgh, PA catalog: 32854) and the Agilent High Sensitivity DNA Bioanalyzer kit (Agilent, Santa Clara, CA, catalog: 5067-4626). Samples had an average distribution between 225 to 275 base pairs. Exome hybridization was performed using the Agilent Human all exon V6 baits (Agilent, Santa Clara, CA, catalog: 5190-8863) using 750 ng of each sample. All samples underwent individual, 24-hour hybridization to the exome baits. Exome capture and post hybridization library prep was performed using Agilent's SureSelect XT HS2 hybridization and blocking reagents (Agilent, Santa Clara, CA, catalog#: G9981A) and fifty ul of MyOne Streptavidin T1 beads (ThermoFisher Scientific, Pittsburgh, PA, catalog #:65602).

**Final library amplification of eleven cycles added the individual index adaptors to the captured libraries.** PCR conditions (98°C 2 min; 98°C 30 sec., 57°C 30 sec., 72°C 1 min (repeat for 11 cycles) and 72°C for 1 min). For all clean-up steps, we used SpriSelect beads (Beckman Coulter catalog# B23318, Indianapolis, IN 46268) and 70% ethanol. Beads were incubated for 5 minutes, washed twice with 70% ethanol, and dried on a 37°C heat block for less than 5 minutes prior to elution. Final library concentration and average library size was determined using the double stranded High Sensitivity Qubit kit (ThermoFisher Scientific, Pittsburgh, PA, catalog#: 32854) and the Agilent High sensitivity DNA Bioanalyzer kit (Agilent, Santa Clara, CA, catalog#:

5067-4626). Final libraries were pooled to 2ng/ul in 130 ul of nuclease free water and shipped for sequencing at Novogene.

**Somatic mutation detection and tumor mutational burden estimation**

After quality control (QC), PE reads were processed to identify somatic mutations (including single nuclear variants, SNP, and small insertions / deletions, indels) following GATK's best practice. Reads were aligned to GRCh38 reference genome (source: GDC) using bwamem with soft-clipping turned on, followed by filtering at mapping quality (MapQ) ≥30, duplicate removal, and base quality score recalibration (BQSR). Somatic mutations were identified using GATK4-MuTect2 with the tumor-normal mode (matched normal tissues from the same patients as control), and the vendor provided target bed file with 100bp padding. Off-target variant calls were removed. High-quality somatic mutations passing MuTect2's intrinsic filters including read orientation filters, oxidative artifact filters, among others (labeled as PASS), and at base quality (BaseQ) ≥20, support reads ≥8, were carried on for annotation by GATK4-Funcotator. Total tumor mutational burden (TMB) was computed by using all predicted protein-coding somatic mutations, including missense SNV, nonsense SNV, frameshift and non-frameshift indels, variants affecting transcriptional or translation starting sites, and variants affecting splice sites, from the target regions on the genome.

**Somatic copy number variation detection**

Somatic copy number variations (CNVs) were identified using the GATK4 somatic CNV workflow, with a panel of normals (PoN) constructed from matched normal tissues from the same patients that the PDOs were derived from. The pipeline includes preprocessing and denoising steps using the PoN as a reference, followed by allelic copy ratio analysis leveraging common germline SNPs (e.g., from dbSNP). For WES data, GATK segments the genome based on read depth and allelic imbalance, requiring at least 5 exons or bins per segment, and estimates copy number per segment after denoising. Resulting CNV segments were further annotated using GISTIC2, which mapped CNVs to gene regions and identified focal and broad alterations.

**Clonality analysis**

Molecular clonality was reconstructed by RETCHER (accessed 01/24/2025) with the original tumor and serial PDO samples from individual patients. RETCHER takes in the allele frequencies of somatic mutations, major and minor copy numbers of somatic CNVs, clusters mutations, determines clonality architecture, and reconstructs phylogenetic trees. To generate the somatic mutation input files for RETCHER, we created a union set of high-quality somatic SNVs from all samples of individual patients including the original tumor and PDOs, and extracted allele fractions at those sites from the alignment BAM files of each sample using GATK4-ASEReadCounter. Time-course subclonal architecture was visualized using fishplot (v0.5.2) and timescape (accessed 01/31/2025).

**Whole transcriptome RNAseq library preparation and sequencing**

Samples were treated with 100 nM lenvatinib (MedChemExpress, cat no. HY-10981, CAS 417716-92-8) for 24 hours. RNA was extracted from frozen samples using the Qiagen RNeasy Mini Kit (cat. no. 74104), and measured on a Nanodrop before library preparation. The RNA was reverse-transcribed to cDNA and amplified using Illumina® Stranded mRNA Prep from Illumina, Inc (San Diego, CA). The library preparation processes such as adenylation, ligation and amplification were performed following the manual provided by the manufacturer[8 9]. The quantity and quality of the libraries were assessed through Bioanalyzer 2100 and Qubit instruments. Paired-end (PE) 75bp reads were generated on AVITI sequencer following the manufacturer's manual[10], with on average 20 million read pairs per sample.

**RNAseq gene set enrichment analysis (GESA)**

Raw FastQ files were processed in the same way as our previous work[18]. In brief, after quality control, reads were pseudoaligned against human reference transcriptome (GRCh38) and Gencode annotation (v28) to estimate transcript abundance using kallisto[20] (v0.48.0), summarized into gene level by tximport (v1.22.0), normalized by TMM method, and $\log_2$ transformed. Lowly expressed genes were removed prior to statistical comparisons. Differentially expressed genes (DEGs) were identified in comparing treated to untreated PDOs (from same patients) using limma voom with precision weights (v3.54.2) with patient id as the blocking factor. Gene set enrichment analysis (GSEA) was performed on DEGs with MSigDB Hallmark 50 gene sets (H50) using R package fGSEA (v1.34.0).

**Drug sensitivity assessment with cisplatin, cetuximab, lenvatinib**

Drug response was evaluated in opaque-walled 96-well spheroid microplates (Corning). At the time of passage, organoids were trypsinized into a single cell suspension. 10,000 cells were seeded per well in a 10 µL drop of 70% BME, then 100 µL of organoid medium was added after the gel solidified. After allowing three days for organoids to form, serial dilutions of drug were added to the wells in 100 µL of medium, bringing the total volume in the well to 200 µL. Untreated wells (media only, not containing drug) were used as a negative control, and 1µm staurosporine was used as a positive control for cell death. After treatment with drug for three days, the readout was performed using the Cell Titer Glo 3D assay (Promega) according to the manufacturer's instructions. Luminescence readings were taken on a Tecan Spark plate reader and normalized to controls (untreated wells = 100% viability, staurosporine treated wells = 0% viability). Drug information is provided in **Table S7**.

**Radiation and Chemoradiation**

Cells were seeded as above into clear U-bottom 96-well plates. After three days of growth, either medium containing drug or medium without drug was added to the wells, as above. One day after adding drug, plates were irradiated in the Precision CellRad+ x-ray irradiator, at 150 kV and 6.25 mA. Each plate received an individual dose of 0, 1, 2, 4, 8, or 16 Gy. Three days after irradiating organoids, the readout was performed as above using the Cell Titer Glo 3D assay.

**Drug treatment, radiation, chemoradiation data analysis**

Drug information is provided in **Table S7**. As a quality control for the organoid drug screening plates, Z'-scores were calculated as previously described[1] using the negative (untreated) controls and positive (staurosporine) controls. A Z'-score of 0.3 or higher was considered to be a good quality plate, and any plate with a Z'-score less than 0.3 was not included in the data shown here. For all samples on drug screening plates, viability was calculated relative to untreated control (100% viability), and background was subtracted using the staurosporine control (0% viability). For irradiated plates, viability for organoids in all plates was calculated using the 0 Gy control plate. IC50 were not calculated for lenvatinib or cetuximab because cell viability did not drop below 50%. For cisplatin screens, IC50 values were calculated in Graphpad Prism (v10).

**Statistical analysis**

Tumor weight between cases that require Nutlin-3 selection or not was compared using Welch two-sample *t*-test. Organoid maximum diameter between normal and tumor PDOs was compared using linear-mixed effect models (LMMs) with PDO type as the fixed effect and slide id as the random effect. DEGs from RNAseq were identified using explicit Bayesian regression models from limma voom. Cell viability changes upon increasing radiation doses were tested using linear mixed-effects models (LMMs), following protocols of our previous work[11]. P-values from multiple comparisons were adjusted using Benjamini-Hochberg false discovery rate (BH-FDR) procedure.

**Data Availability**

The de-identified WES data generated in this study will be deposited to NCBI GEO repository. Processed data relevant to the study are included in the article or uploaded as supplementary information. Additional data is provided on GitHub repository: https://github.com/HCC-data-sciences-pub/HNSCC-PDO-integrative-analysis. This study utilized open-source tools as described in **Supplementary Methods**. Other data will be provided upon request from the corresponding authors.

### Supplementary Figures

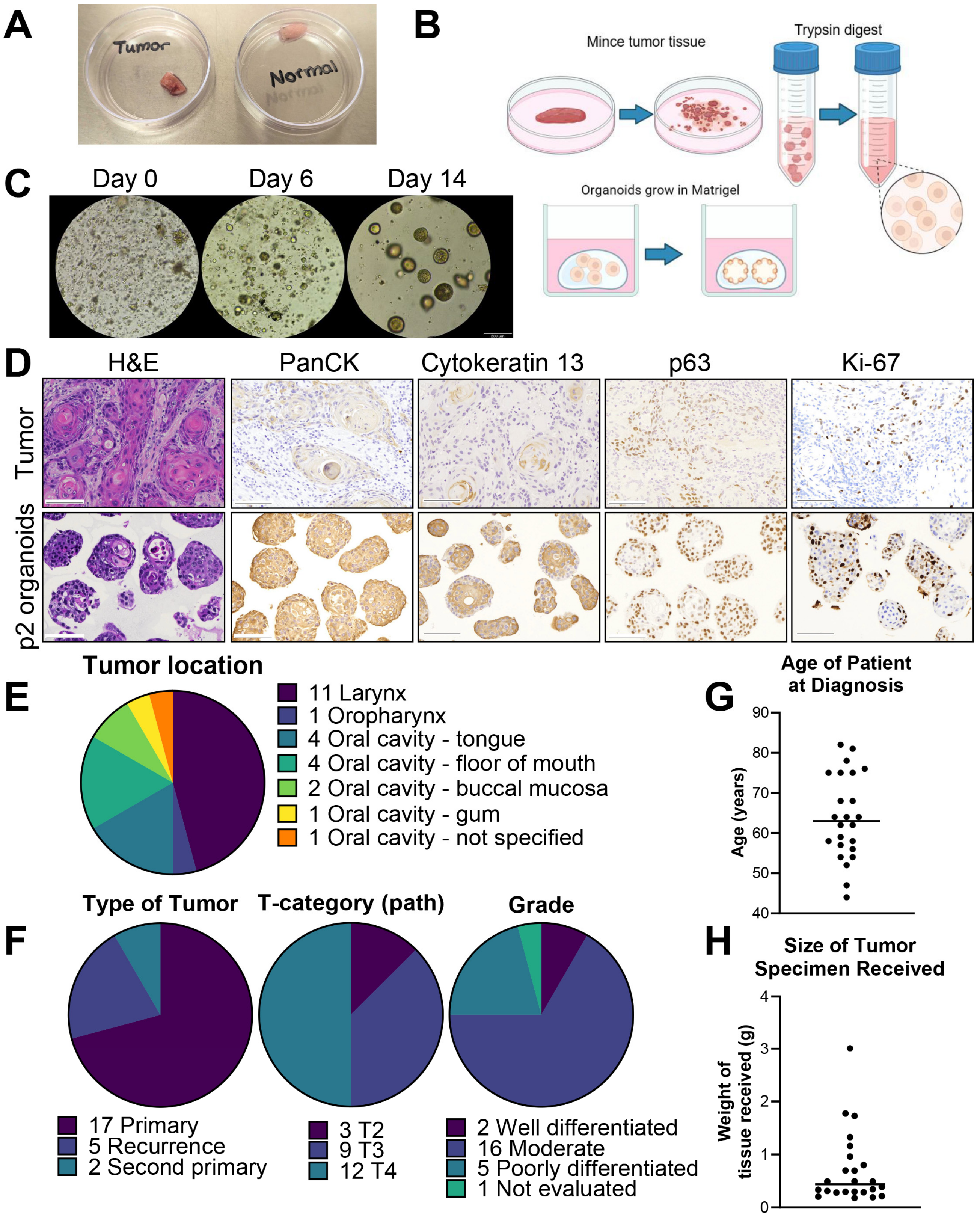


**Supplementary Figure 1. Overview of PDO development workflow, morphological features, and clinical cohort characteristics.** (**A**) Representative photograph of fresh tumor and normal tissue samples received for PDO culture. (**B**) Schematic of the organoid development workflow. Tumor tissue is minced, enzymatically digested with trypsin, and embedded in Matrigel for 3D culture. (**C**) Brightfield images showing PDO growth over time. Cells transition from dispersed single cells on Day 0 to mature spherical organoids by Day 14. (**D**) Histologic and immunohistochemical (IHC) comparison of original tumor tissue and passage 2 (p2) PDOs. Tumor and organoids were stained with H&E, PanCK, p63, Cytokeratin 13, and Ki-67. (**E**) Pie chart showing distribution of tumor anatomical locations within the cohort. (**F**) Clinical and pathological features of the PDO cohort, including tumor type, T-category (pathologic), and grade. (**G**) Age distribution of patients at time of diagnosis. (**H**) Distribution of tissue specimen weights received for PDO generation.

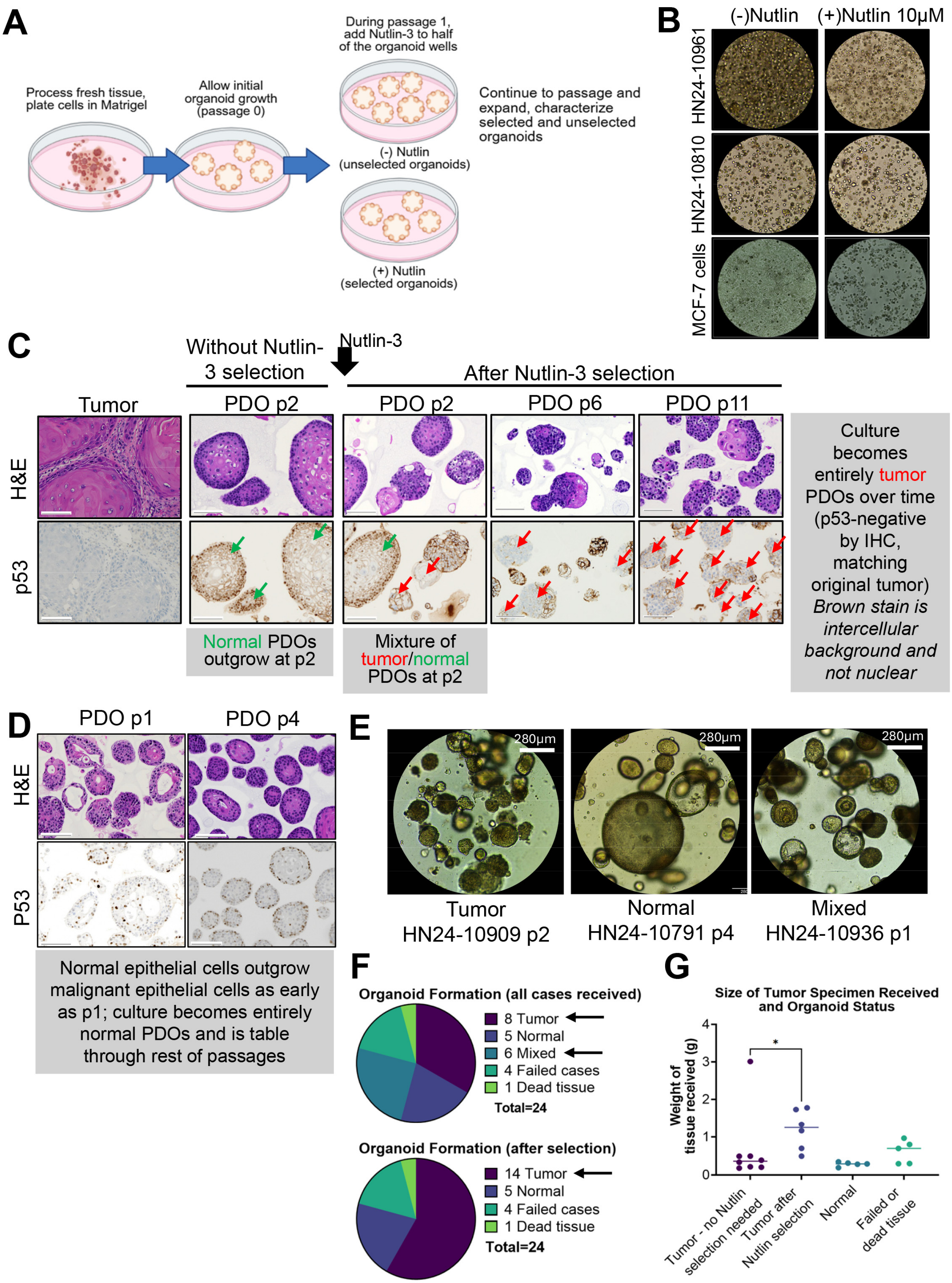


**Supplementary Figure 2. Nutlin-3 selection enriches tumor PDOs in mixed cultures.** (**A**) Schematic of the Nutlin-3 selection strategy. Following initial organoid formation (passage 0), Nutlin-3 (10 μM) was added to half of the wells during passage 1. PDOs were expanded and characterized in Nutlin-treated and untreated conditions. (**B**) Representative brightfield images of organoid cultures treated with Nutlin-3, including mixed tumor and normal PDOs (top), all tumor PDOs (middle) and control MCF-7 cells (bottom). (**C**) H&E and p53 IHC of tumor tissue and matched PDOs with and without Nutlin-3 selection. Without selection, organoid cultures showed wild-type p53 staining, consistent with normal epithelial cell outgrowth. After selection, p53-positive malignant PDOs were enriched (red arrows). (**D**) Serial passages of PDOs derived from a normal outgrowth case, showing how normal epithelial cells overtake cultures over time. (**E**) Brightfield microscopy images of tumor, normal, and mixed PDO cultures. (**F**) Pie charts summarizing PDO types from 24 total cases, before and after Nutlin-3 selection. Selection increased the number of tumor PDOs and reduced mixed or normal outgrowth cases. (**G**) Tumor specimen weights grouped by final PDO status. Tumor PDOs without Nutlin selection tend to derive from smaller tissue specimens compared to those requiring selection or resulting in normal PDOs. Two-sided Welch two-sample *t*-test was used in **G**. Denotation: * $P < 0.05$.

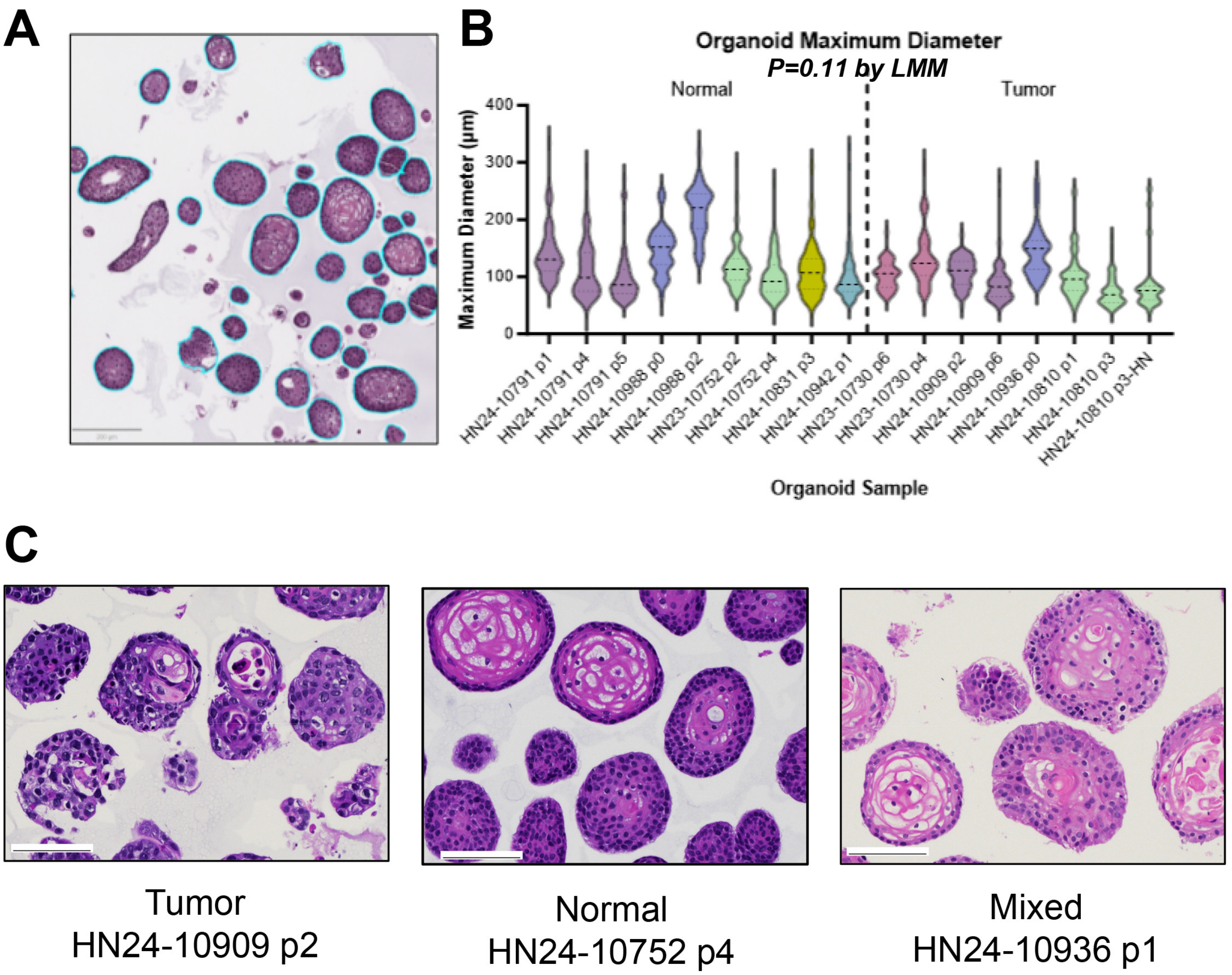


**Supplementary Figure 3. Morphometric and histologic features of tumor and normal PDOs.** (**A**) Representative H&E image of a PDO section with individual organoid boundaries outlined in blue. Organoid morphology was manually annotated from high-resolution images. (**B**) Violin plot showing maximum diameters of individual organoids across PDO samples, grouped by type (tumor or normal). *P* = 0.11 was shown comparing the two groups of PDOs. (**C**) H&E images of tumor, normal, and mixed PDO cultures. Linear mixed-effects models (LMM) were used in **A**, with PDO type as the fixed effect and slide id as the random effect.

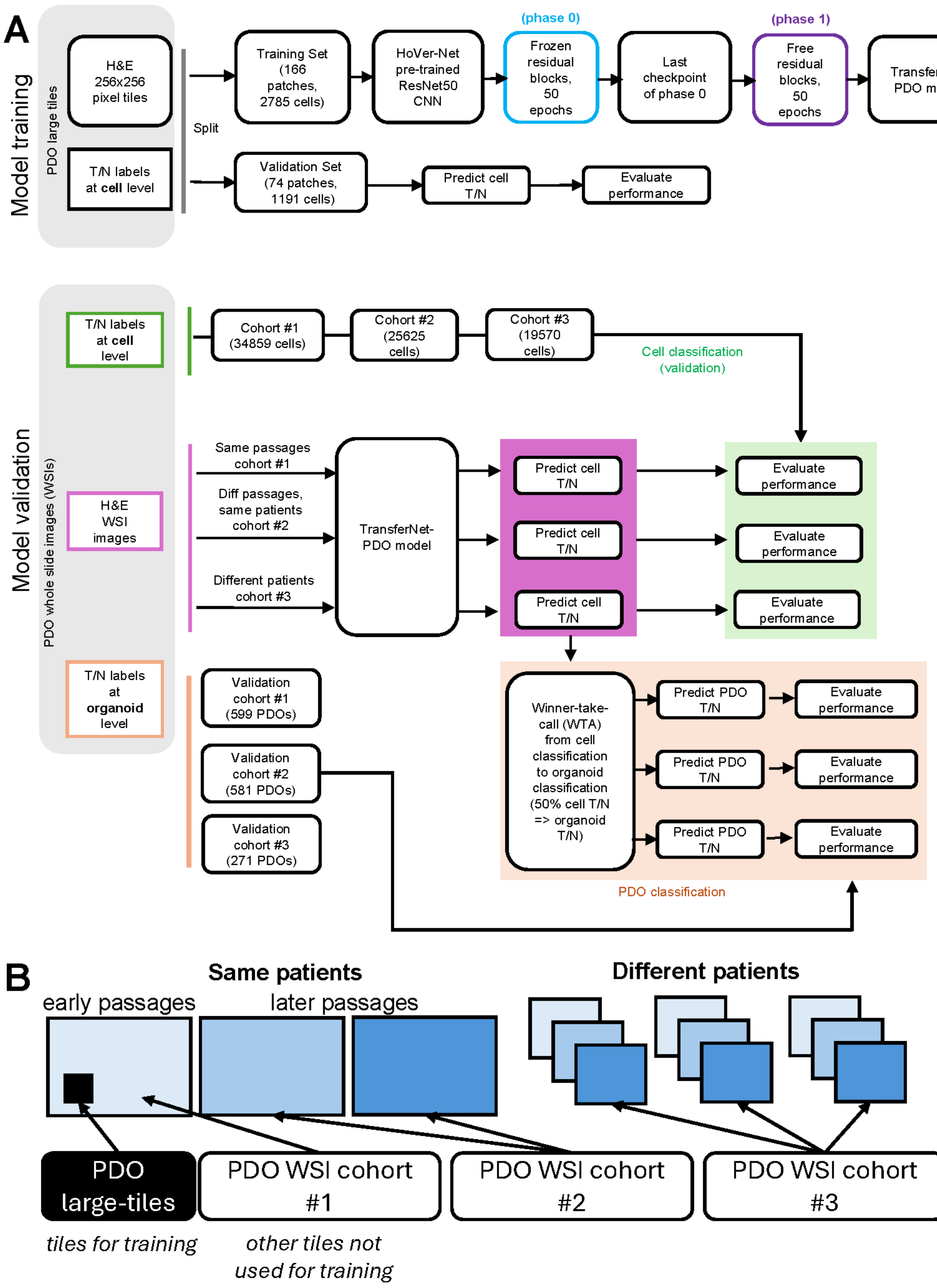


**Supplementary Figure 4. TransferNet-PDO development workflow**. (**A**) Schematic of the TransferNet-PDO training and validation pipeline. (**top**) For model training, H&E-stained 256×256-pixel tiles with cell-level tumor/normal (T/N) annotations were split into training (2,785 cells) and validation (1,191 cells) sets. A Hover-

Net CNN ResNet50 pre-trained model was used for feature extraction. During phase 0, residual blocks were frozen, and classification weights were updated. In phase 1, all layers were unfrozen for full fine-tuning over 50 epochs. The final model was selected based on validation performance. (**bottom**) For independent validation of the final model selected from the training process, cell-level predictions were tested on three cohorts of WSIs: cohort 1 (same images as training, different regions), cohort 2 (different passages from same patients), and cohort 3 (different patients). Model performance was evaluated both at the single-cell level and aggregated to the PDO level using a winner-take-all (WTA) strategy (i.e., majority of cells define organoid label). (**B**) Schematic of WSI cohort structure for independent validation. Training tiles (black solid box) were selected from early passage WSIs. Validation cohorts consisted of non-overlapping regions from the same slides (cohort 1), later passage WSIs from the same patients (cohort 2), and WSIs from entirely different patients (cohort 3).

(1) Nucleated Cell Detection Cohort (large tiles)

**Cell Detection — Training**

| Actual \ Predicted | Cell is true | Cell is not true | total |
|---|---|---|---|
| Cell is true | 2785 | 250 | 3035 |
| Cell is not true | 276 | NA | NA |

**Cell Detection — Validation**

| Actual \ Predicted | Cell is true | Cell is not true | total |
|---|---|---|---|
| Cell is true | 1191 | 110 | 1301 |
| Cell is not true | 112 | NA | NA |

**Legend**

| Actual \ Predicted | | |
|---|---|---|
| | TP | FN |
| | FP | TN |

(2) **Cell Classification** (tumor/normal, tumor as positive label)

Cell Training Cohort (large tiles)

**Cell Classification — Training (2785 cells)**

| Actual \ Predicted | Cell is tumor | Cell is normal | total |
|---|---|---|---|
| Cell is tumor | 1294 | 33 | 1327 |
| Cell is normal | 76 | 1382 | 1458 |

**Cell Classification — Validation (1191 cells)**

| Actual \ Predicted | Cell is tumor | Cell is normal | total |
|---|---|---|---|
| Cell is tumor | 358 | 10 | 368 |
| Cell is normal | 55 | 768 | 823 |

Cell Validation Cohorts #1-#3 (WSIs)

**Cell Classification — Cohort #1 same passages**

| Actual \ Predicted | Cell is tumor | Cell is normal | total |
|---|---|---|---|
| Cell is tumor | 12217 | 281 | 12498 |
| Cell is normal | 4686 | 17675 | 22361 |

**Cell Classification — Cohort #2 diff passages, same patients**

| Actual \ Predicted | Cell is tumor | Cell is normal | total |
|---|---|---|---|
| Cell is tumor | 6542 | 1824 | 8366 |
| Cell is normal | 4620 | 12639 | 17259 |

**Cell Classification — Cohort #3 different patients**

| Actual \ Predicted | Cell is tumor | Cell is normal | total |
|---|---|---|---|
| Cell is tumor | 4266 | 1252 | 5518 |
| Cell is normal | 4767 | 9285 | 14052 |

(3) **PDO Classification** (tumor/normal, tumor as positive label)

PDO Prediction Cohorts #1-#3 (WSIs)

**PDO Classification — Cohort #1 same passages**

| Actual \ Predicted | PDO is tumor | PDO is normal | total |
|---|---|---|---|
| PDO is tumor | 268 | 32 | 300 |
| PDO is normal | 1 | 298 | 299 |

**PDO Classification — Cohort #2 diff passages, same patients**

| Actual \ Predicted | PDO is tumor | PDO is normal | total |
|---|---|---|---|
| PDO is tumor | 262 | 72 | 334 |
| PDO is normal | 35 | 212 | 247 |

**PDO Classification — Cohort #3 different patients**

| Actual \ Predicted | PDO is tumor | PDO is normal | total |
|---|---|---|---|
| PDO is tumor | 60 | 14 | 74 |
| PDO is normal | 45 | 152 | 197 |

**Supplementary Figure 5. Confusion matrices summarizing TransferNet-PDO performance for cell and PDO classification.** (1) Confusion matrices for cell detection and classification in training & validation sets from largetile PDO images. (**top**) Cell detection performance (nucleated cell is present or not). (**bottom**) Cell classification performance (cell-level tumor *versus* normal). (2) Cell classification performance across three independent whole slide image (WSI) validation cohorts. Cohort #1: same slides as training but different regions; Cohort #2: different passages from the same patients; Cohort #3: entirely different patients. (3) PDO-level classification performance derived from cell-level predictions by winner-take-all (WTA). PDOs were labeled as tumor or normal based on majority of constituent cell classifications. True positive (TP), false positive (FP), true negative (TN), and false negative (FN) are color-coded across all panels for interpretability.

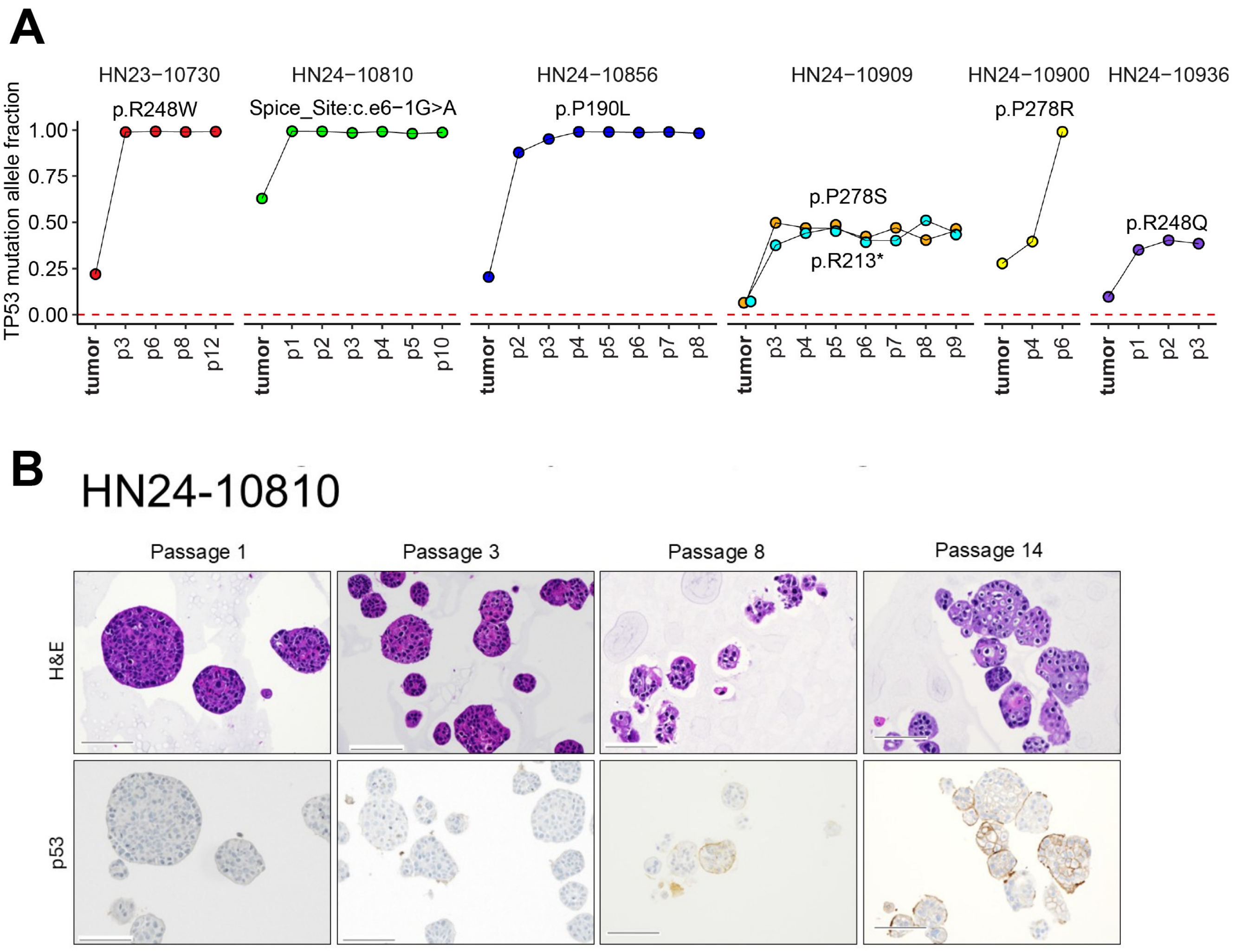


**Supplementary Figure 6. Longitudinal analysis of *TP53* mutation retention and p53 protein expression across PDO passages.** (**A**) *TP53* mutation allele fractions measured by whole-exome sequencing (WES) in tumors and corresponding PDOs across serial passages. Each line represents a unique patient case. Mutant allele fractions remained stable across passages, confirming retention of driver *TP53* mutations in PDOs. Cases with multiple TP53 mutations (e.g., HN24-10909) are indicated. Color indicates individual variant. (**B**) Representative H&E and p53 IHC staining of PDOs from patient HN24-10810 (p53-negative mutant) across passages 1, 3, 8, and 14. *TP53* mutation (splice site c.6-1G>A) was consistently retained and potentially resulted in truncated protein due to splice site disruption. This was further validated by absence of protein by IHC. Brown stain was intercellular background and not nuclear staining. Scale bars = 100 μm.

### Supplementary Tables 1-9 Titles

**Supplementary Table 1. Antibodies used for IHC.**

**Supplementary Table 2. Metadata of samples used for whole exome sequencing (WES).**

**Supplementary Table 3. Somatic mutations in known HNSCC cancer genes. Coding variants are shown.**

**Supplementary Table 4. Somatic mutation landscape of HNSCC PDOs and original tumors. Both coding and non-coding variants are shown.**

**Supplementary Table 5. Somatic CNV landscape of HNSCC PDOs and original tumors.**

**Supplementary Table 6. List of somatic mutations delineating the clonal architecture of HNSCC PDOs and original tumors.**

**Supplementary Table 7. Drugs used for drug screening.**

**Supplementary Table 8. IC50 of Cisplatin varies between HNSCC PDOs from different patients.**

**Supplementary Table 9. Gene set enrichment analysis (GESA) results of RNAseq gene expression from lenvatinib treatment on HNSCC PDOs.Supplementary Table 10. Assessment of drug sensitivity in combination with radiation in tumor PDOs.**
